# Random Sanitization in DNA information storage using CRISPR-Cas12a

**DOI:** 10.1101/2024.08.04.606549

**Authors:** Hongyu Shen, Zhi Weng, Haipei Zhao, Haitao Song, Fei Wang, Chunhai Fan, Ping Song

**Affiliations:** School of Biomedical Engineering, The International Peace Maternity and Child Health Hospital, Zhangjiang Institute for Advanced Study and National Center for Translational Medicine, Shanghai Jiao Tong University, Shanghai 200240, China; The Institute of Artificial Intelligence and National Center for Translational Medicine, Shanghai Jiao Tong University, Shanghai 200240, China; School of Chemistry and Chemical Engineering, New Cornerstone Science Laboratory, Frontiers Science Center for Transformative Molecules, Zhangjiang Institute for Advanced Study and National Center for Translational Medicine, Shanghai Jiao Tong University, Shanghai 200240, China

## Abstract

DNA information storage provides an excellent solution for metadata storage due to its high density, programmability, and long-term stability. However, current research in DNA storage primarily focuses on the processes of storing and reading data, lacking comprehensive solutions for the secure metadata wiping. Herein, we present a method of random sanitization in DNA information storage using CRISPR-Cas12a (RSDISC) based on precise control of the thermodynamic energy of primer-template hybridization. We utilize the collateral cleavage (trans-activity) of single-stranded DNA (ssDNA) by CRISPR-Cas12a to achieve selective sanitization of files in metadata. This method enables ssDNA degradation with different GC content, lengths, and secondary structures to achieve a sanitization efficiency up to 99.9% for 28,258 oligonucleotides in DNA storage within one round. We demonstrate that the number of erasable files could reach 10^11.7^ based on a model of primer-template hybridization efficiency. Overall, RSDISC provides a random sanitization approach to set the foundation of information encryption, file classification, memory deallocation and accurate reading in DNA data storage.

## Introduction

With the rapid development of information technologies such as the Internet and artificial intelligence, the global volume of data is experiencing explosive growth, and it is expected to reach 175 ZB by 2025^1^. DNA storage, as an emerging information storage technology, has become an ideal solution to address the challenges of big data storage due to its high storage density, programmability, long-term stability, and low energy consumption^2,3^. Current research primarily focuses on encoding and decoding methods^4^, as well as the storage and presentation of multimodal data, with an emphasis on optimizing the storage process and constructing random access systems^5^. However, the importance of data security remains insufficiently addressed^6^. Nevertheless, data security is a critical aspect in ensuring the continuous and effective development of DNA storage technology and is a significant component of information management, particularly in metadata storage^7^.

In the field of information technology, many studies focus on designing highly secure data deletion systems to protect sensitive information^8,9^. For example, the Gutmann algorithm is used for data deletion from magnetic and solid-state storage media^10^, the DoD 5220.22-M standard employs 3-pass or 7-pass overwriting techniques^11^, and the NIST 800-88 guidelines are used for media sanitization^12,13^. In recent years, with the surge in stored data volume, developing fast and effective deletion methods has become increasingly important^14^. Kim et al proposed a method that encodes DNA into a metastable aqueous solution, enabling rapid information deletion through a simple heating process^15^. Although similar approaches achieve selective information deletion, most deletion systems are based on DNA computation or specific biomolecular reaction patterns and do not achieve a complete deletion of data^16-18^. Instead, they merely temporarily obscure the information, resulting in a risk of data breach. Therefore, developing a simple and effective method for random sanitization of storage media to prevent leakage of personal privacy or national secrets is of great importance^19^.

To address this issue, we present random sanitization in DNA information storage using CRISPR-Cas12a (clustered regularly interspaced short palindromic repeats– CRISPR-associated protein) (RSDISC), a selective erasure method with oligonucleotide sanitization efficiency up to 99.9% and specificity up to 99.5%. For each single-stranded DNA (ssDNA) in the pool, we convert the single strands into double strands that we plan to retain, and then the activated Cas12a complex could cleave the residual ssDNA^20^. Our method can cleave essentially any ssDNA with varied GC contents (30%~80%), varied lengths, and secondary structure modifications. Furthermore, because the maximum number of erasable files depends on the hybridization efficiency between primers and templates, it is possible to wipe 10^11.7^ files of up to 10^7^ GB with 100% file sanitization efficiency and 100% specificity. As proof of concept of potential applications, we adapted the RSDISC method to achieving information encryption through permanent erasure. And by completely erasing unnecessary information, it could deallocate memory to alleviate the memory pressure caused by the data surge. On the other hand, the technology may help advance file classification and reduce non-specific hybridization during sequencing.

## Results

### Overview of Selective File Sanitization Based on CRISPR-Cas12a

Existing random deletion methods in the storage rely on random access systems, which only temporarily hide deleted data while the information itself remains intact in the system^21^. To achieve complete information erasure, we have developed a random sanitization method in DNA information storage of RSDISC. This method can thoroughly erase non-target files while preserving specific target files, enabling metadata encryption and memory deallocation (Figure 1a, b and Supplementary Figure S1). Our erasure system involves three key steps: (1) Data encoding: File information is encoded into binary digits and then further translated into the quaternary code of DNA bases (Supplementary Figure S2). (2) Input: Synthetic DNA sequences containing the encoded information are incorporated into the DNA storage repository (Supplementary Figure S3). (3) Metadata sanitization: Target files are protected by amplifying the targeted DNA with reverse primers (RPs), while all other unprotected files are erased, resulting in partial memory deallocation (Figure 1b, Supplementary Figure S4).

**Fig. 1.**
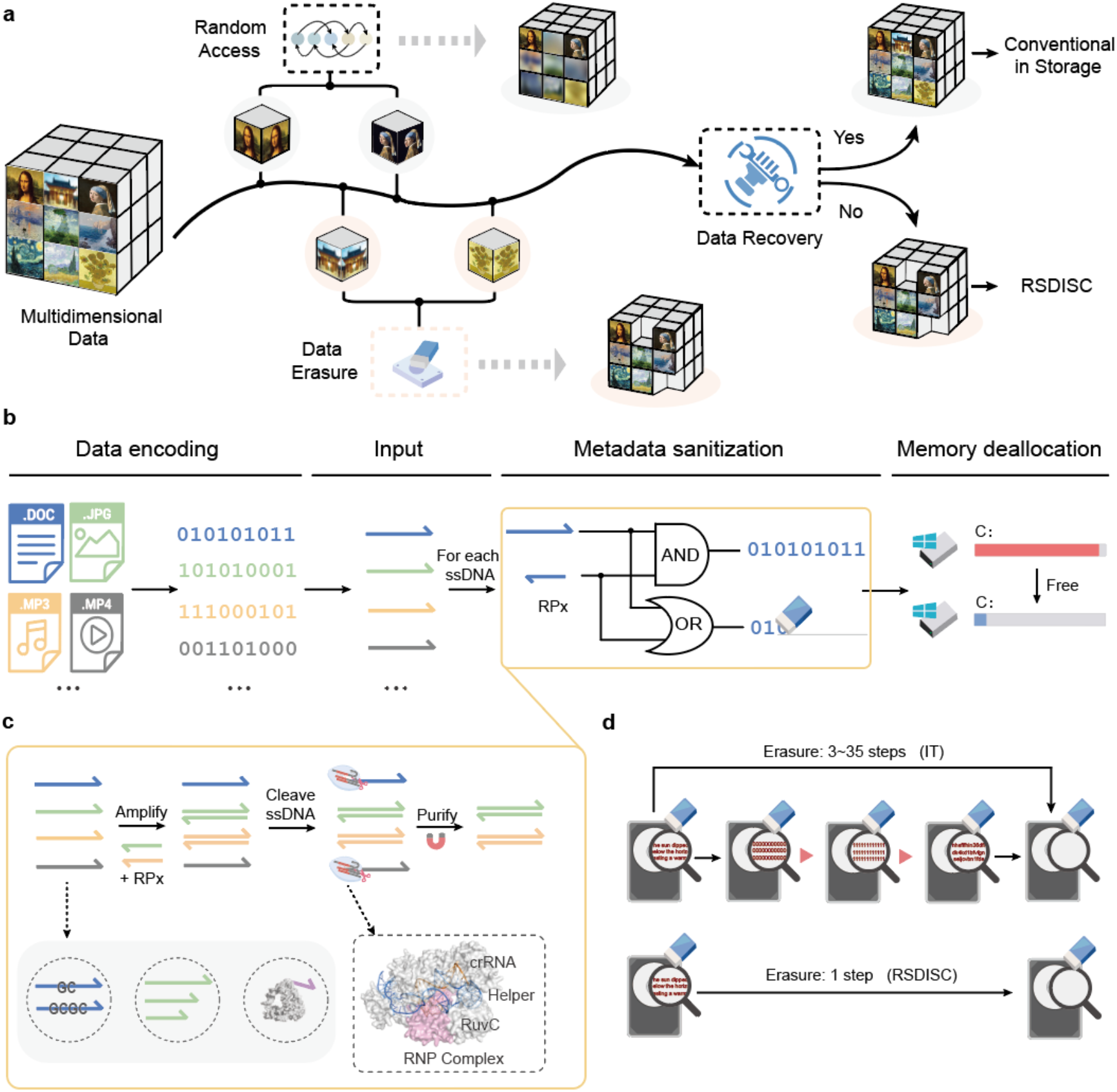
Schematic workflow of RSDISC. **a**, Illustration of the difference between random access and data erasure in DNA information storage. **b**, Complete framework of a random sanitization system in DNA data storage, resulting in information encryption and memory deallocation. **c**, Implementation of selective cleavage of single-stranded DNA (ssDNA) in RSDISC. This assay employs primers to amplify and preserve the targeted single strands while utilizing the trans-cleavage activity of CRISPR-Cas12a to cleave the remaining single strands. Meanwhile, this system can cleave any type of single strand. **d**, A comparison of the RSDISC assay and the IT (information technology) assay. The IT assay requires 3 to 35 steps to delete information, whereas the RSDISC assay achieves information erasure in a single step. IT assay: DoD 5220.22-M and Gutmann algorithm.

In the specific metadata selective erasure process, we first prepared Helper-DNA that targeted binding with CRISPR RNA (crRNA) to form a complex with crRNA-Cas12a, which activated the Cas12a’s trans-cleavage activity. Next, we amplified the targeted ssDNA into a double-stranded DNA (dsDNA) and used the activated ribonucleoprotein (RNP) complex to cleave the remaining single strands^20^ (Figure 1c). To assess the universal applicability of our system to different types of single-stranded substrates, we demonstrated its cleavage capability against various types of ssDNA, including those with varying GC content, lengths, and complex secondary structure modifications.

To address the lack of a method for absolute data erasure in the current DNA storage field, we have developed the RSDISC technology, which achieves permanent information erasure in a single step, with the erased data being irrecoverable by any means (Figure 1d). To validate the feasibility of this technology in practical storage, we demonstrated that our system achieves up to 99.9% oligonucleotides sanitization efficiency and 99.5% oligonucleotides sanitization specificity in a storage pool containing 28,258 ssDNA oligonucleotides. Additionally, our technology can be used for information protection, memory deallocation, information classification, and improving sequencing accuracy by reducing non-specific hybridization during library preparation.

### Cleavage of Different Types of Templates in a Single System

To investigate the effectiveness of this method in a simple system, we initially tested its cleavage efficiency using a single type of ssDNA. Before comparing the cleavage capabilities for different substrates, we optimized the reaction conditions for CRISPR-Cas12a^22^ (Supplementary Figures 5 and 6). To activate the trans-cleavage activity of Cas12a, we used RPs with an adapter (containing a sequence complementary to crRNA) to amplify the substrate (pre-Helper) and prepare single-ended Helper-DNA (S-Helper) system with targeted crRNA. This system was then incubated with crRNA-Cas12a to form an RNP complex capable of cleaving any collateral ssDNA (Figure 2a). To test the system’s cleavage capability on long ssDNA (>100 nt), we found that the cleavage efficiency for 130 nt ssDNA reached 99% at the plateau phase (Figure 2b and Supplementary Figures 7 and 8). In addition to conventional substrates, we also explored the cleavage effectiveness of the system on ssDNA oligonucleotides with high GC contents, different lengths, and complex secondary structures. We found that our system could almost cleave essential any ssDNA regardless of GC content, length, or complex structure, with cleavage efficiency reaching up to 99% (Figure 2c).

**Fig. 2.**
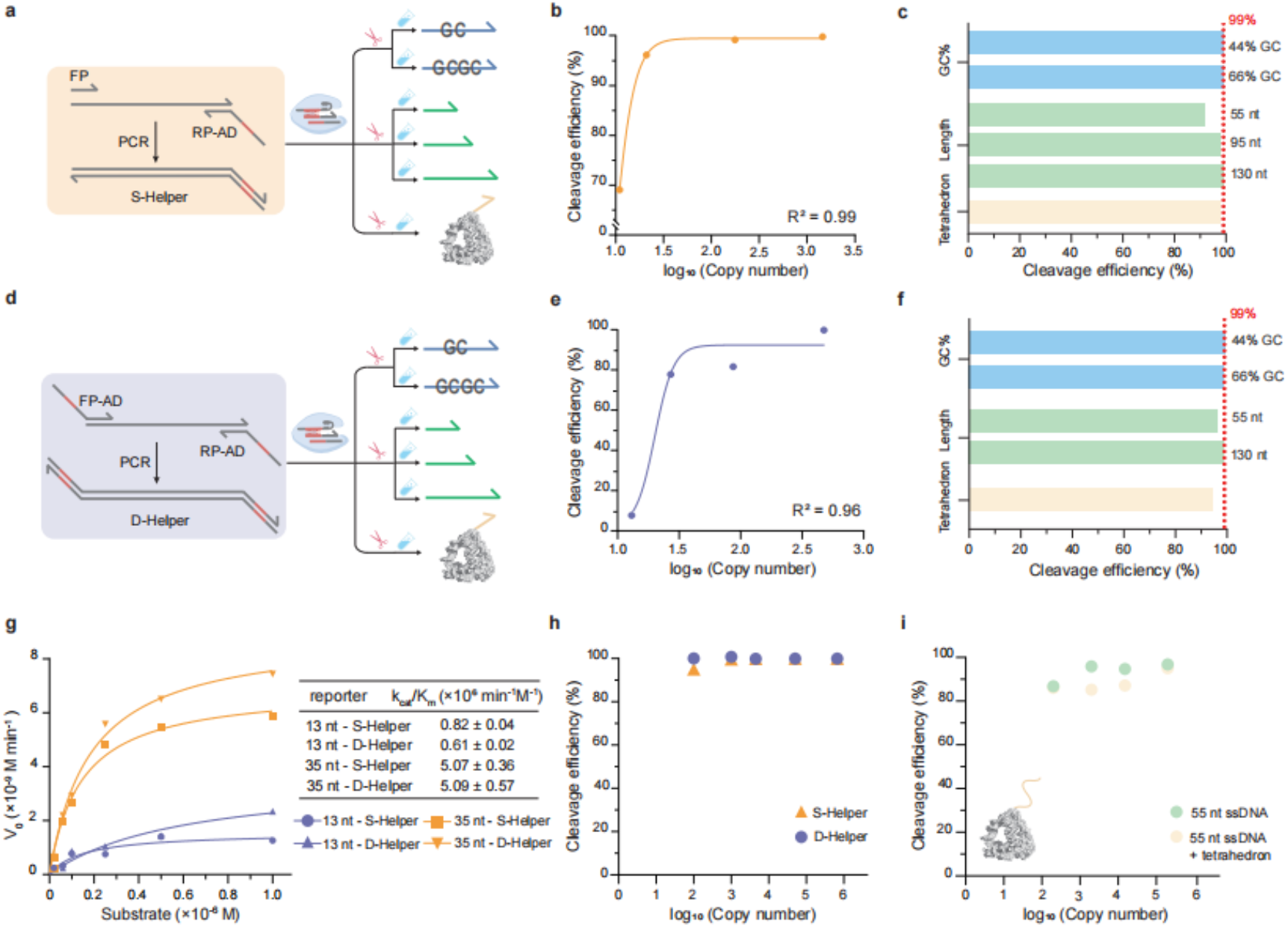
Cleavage performance of single-end and double-end systems to activate Cas12a in a single system. **a**, Activation of the single-end Helper-DNA (S-Helper) system of Cas12a cleaves ssDNA with varying GC content, different lengths, and three-dimensional structures. **b**, Calibrated curve of 130 nt ssDNA’s cleavage efficiency by Cas12a activated with S-Helper system. The detection limit for single-strand cleavage in this system is 11 copies. **c**, Comparison the cleavage efficiency of ssDNA with different GC content, lengths, and secondary structure modifications after activation of Cas12a with S-Helper system. **d**, Activation of the double-end Helper-DNA (D-Helper) system of Cas12a cleaves single-stranded DNA with varying GC content, different lengths, and three-dimensional structures. **e**, Calibrated curve of 130 nt ssDNA’s cleavage efficiency by Cas12a activated with D-Helper system. The detection limit for single-strand cleavage in this system is 13 copies. **f**, Comparison the cleavage efficiency of ssDNA with different GC content, lengths, and secondary structure modifications after activation of Cas12a with D-Helper system. **g**, Comparison of the kinetic parameters of Cas12a in S-Helper and D-Helper systems using 13 or 35 nt reporters. 13/35 nt-S(D)-Helper, cleave 13/35 nt reporter with S(D)-Helper system; measured k_cat_/K_m_ values reported as mean ± SD, where n = 3 Michaelis Menten fits. V_0_, rate of catalysis. Substrate, 13 nt or 35 nt reporters. **h**, Comparison of cleaving different copies of ssDNA substrate using S-Helper *or* D-Helper systems to activate the trans-activity of Cas12a. **i**, Performance comparison between the cleavage effectiveness of ssDNA and ssDNA with tetrahedron modification using D-Helper system to activate Cas12a. The tetrahedron has a typical edge length of ~6 nm (17 base pairs for each edge).

Furthermore, we hypothesized that increasing the number of the activated RNP complex per unit volume might enhance the chances of ssDNA interacting with Cas12a, potentially improving cleavage efficiency. Therefore, we planned to use forward primers (FPs) with an adapter along with RPs to simultaneously amplify the pre-Helper and synthesize double-ended Helper-DNA (D-Helper) system with targeted crRNA (Figure 2d, Supplementary Figure S8). Experimental results showed that the D-Helper system incubating with Cas12a achieved a cleavage efficiency of 92% for 130 nt ssDNA at the plateau phase (Figure 2e, Supplementary Figure S9). Similarly, we found that the GC content, length, and complex structural modification of the ssDNA did not significantly affect the system’s cleavage capability (Figure 2f). Overall, we demonstrated that both S-Helper and D-Helper systems can efficiently cleave ssDNA with varied GC content (30% - 80%), varied length, and modified secondary structures. To compare whether there are significant differences in the kinetics of cleavage reactions between S-Helper and D-Helper systems that activating Cas12a, we calculated the catalytic efficiency (k_cat_/K_m_) of 13 nt and 35 nt fluorophore-quenched (FQ) labeled ssDNA reporter substrates (Supplementary Figures 10, 11 and 12) in both systems^20,23-25^. The experimental results indicated that, regardless of whether 13 nt or 35 nt reporter is used, the catalytic efficiency of the same substrate in the S-Helper and D-Helper systems is almost identical (Figure 2g and Supplementary Figure 13). And by comparing the catalytic efficiency of different substrates in the same system, we found that the slight differences in efficiency could be attributed to the different fluorophore groups attached to the reporters: the 13 nt reporter was labeled with VIC, while the 35 nt reporter was labeled with Cy5^26^. Further comparison of the cleavage efficiency for high concentrations of the same ssDNA in both systems showed that both achieved a cleavage efficiency of 99% (Figure 2h), which is consistent with previous experimental results.

Although our previous experiments demonstrated that our system performs well on single strands with complex secondary structure modifications, we sought to further investigate whether spatial hindrance effects due to secondary structures could obstruct the cleavage efficiency (Supplementary Figure 14). Therefore, we attached a DNA tetrahedral framework with a side length of ~6 nm (each edge containing 17 bases) to the end of an exposed 55 nt ssDNA^27,28^. By comparing their cleavage efficiencies, we observed that the efficiency difference between samples ranged from 0.7% to 10.6%. These suggested that spatial hindrance effects caused by complex secondary structures may moderately impact the cleavage efficiency of the single strands itself (Figure 2i).

### Selective Cleavage of Templates in a Multiplex System

Given that our two systems demonstrate excellent cleavage efficiency in single systems, we chose the D-Helper system activating Cas12a to further investigate its ability to selectively cleave ssDNA in complex multi-template systems with template interactions (Figure 3a). In a multi-template system with single strands of varying GC content, we found that the cleavage efficiency for all templates was greater than 90% (Figure 3b). Additionally, although previous studies have found that Cas12a may show preference for short poly A or T reporters^29,30^, we observed that adding a 10 nt poly A/T sequence to the 5’ or 3’ end of a 130 nt ssDNA did not significantly enhance its cleavage efficiency (Figure 3b and Supplementary Figure 15). To investigate whether the sequence of crRNA would affect cleavage efficiency^31^, we designed two different hybridization sequences of crRNA to activate Cas12a. The results showed that both sequences achieved cleavage efficiencies of up to 99% (Figure 3c). Next, we examined the efficiency of ssDNA selective retention and cleavage in a five-template system (Figure 3d). To verify the specificity of this system for ssDNA retention, we selected seq4 and seq5 as targeted protection sequences. The experimental results showed that both seq4 and seq5 were well retained, with non-specific cleavage efficiencies of approximately 5% for each. All non-target ssDNA oligonucleotides were completely cleaved (with cleavage efficiencies reaching 99%), indicating that our method could efficiently and specifically cleave ssDNA. Furthermore, to demonstrate that the system does not affect the sequence of protected ssDNA, we performed Sanger sequencing on the seq5 products treated with Cas12a. The sequencing results showed that the base sequence of seq5 remained completely unchanged before and after Cas12a treatment (Figure 3e).

**Fig. 3.**
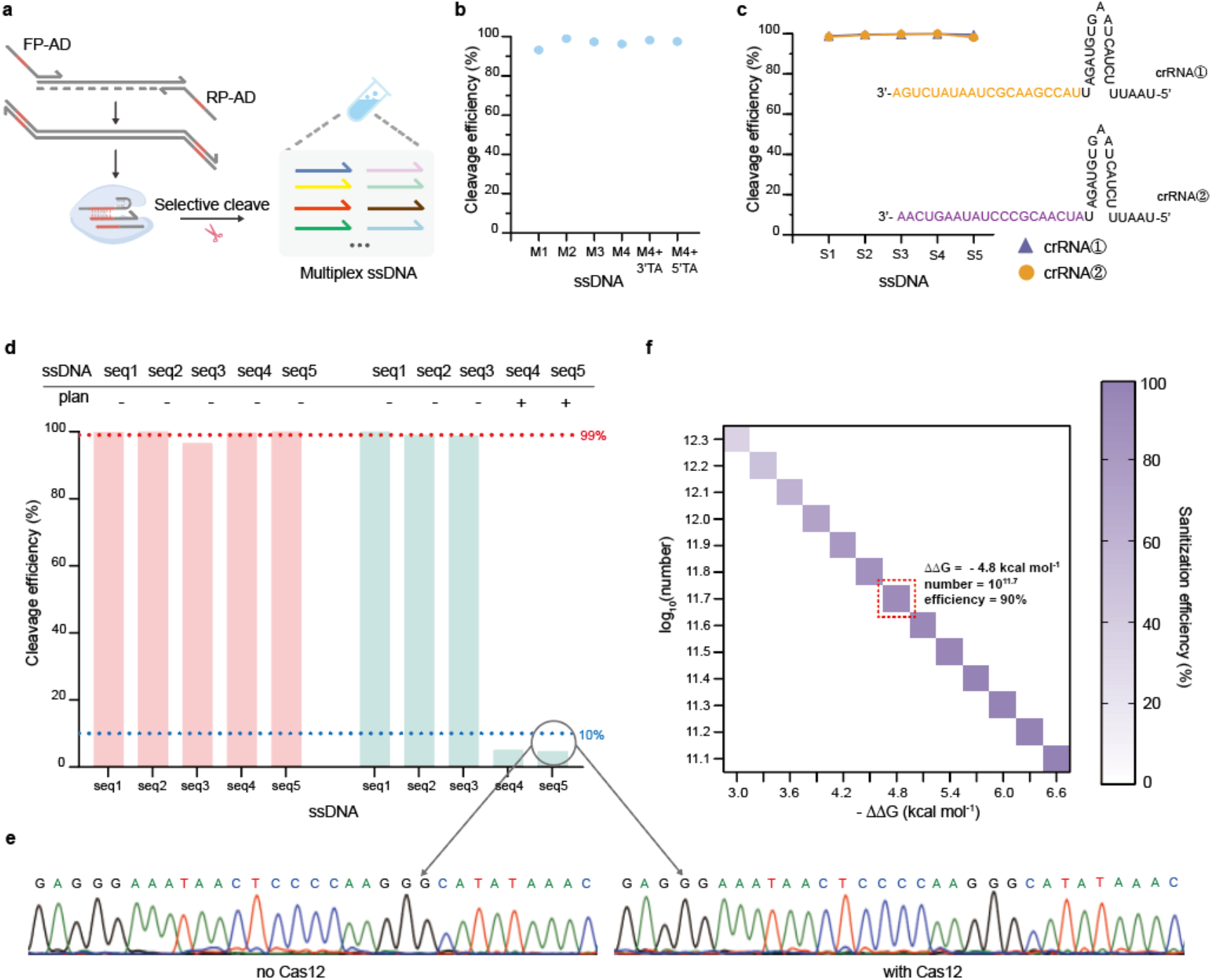
Selective cleavage of ssDNA in a multiplex system. **a**, The assay that achieves selective cleavage of ssDNA in a mixed system using D-Helper system activating Cas12a. **b**, The cleavage effect of ssDNA with different GC contents at the same substrate concentration. M4 + 3’ TA and M4+5’ TA refer to the addition of the ‘TTAATAATTT’ sequence at the 3’ and 5’ end of the M4 template, respectively; **c**, Cleavage activity of different sequences in CRISPR RNA (crRNA) that is complementary to the sequence of Helper-DNA to activate Cas12a. **d**, The effect of selectively retaining different numbers of ssDNA in a multiplex system. **e**, Comparison of Sanger sequencing results of the target ssDNA in with/no Cas12a-treated samples. **f**, Simulated the impact of hybridization efficiency on the maximum number of erasable files. ΔΔG corresponds to the number of base mismatches during hybridization between primers and templates.

To evaluate whether our method is compatible with the current highest storage capacity, we simulated the maximum number of files that can be erased using our method. Our previous experiments have demonstrated that the maximum number of files is determined by the amplification efficiency of primers and templates during the polymerase chain reaction (PCR) process, rather than the concentration of Helper-DNA or activated Cas12a. Therefore, based on hybridization efficiency, we simulated that the maximum number of files that can be erased is approximately 10^11.7^ (satisfying an erasure efficiency >90%), equivalent to 10^7^ GB of data (Figure 3f).

### Selective Sanitization of Files in DNA Storage

To access whether our method can be practically applied in complex storage systems, we investigated its effectiveness in selectively erasing files in a DNA storage pool. For a DNA storage pool containing 28,258 different sequences, we constructed a library containing information from seven files, where the DNA templates encoding the same files share a common address sequence at both ends. Among the seven files, there are three image files and four text files, with the following numbers of sequences encoded for each file: MonaLisa.bmp (ML, 13,490 oligonucleotides), SJTU.bmp (SJTU, 3,698 oligonucleotides), FlightRoute.bmp (FR, 5,454 oligonucleotides), TheAnalectsOfConfucius.txt (AC, 3,593 oligonucleotides), ThreeCharacterClassic.txt (TCC, 315 oligonucleotides), TaoTeChing.txt (TTC, 1,054 oligonucleotides), and IHaveADream.txt (IHAD, 654 oligonucleotides). In the selective erasure step, we used RPs to amplify the ssDNA encoding the targeted information into double strands for retention. Then, we employed activated RNP complex to cleave the remaining single strands, achieving selective erasure of information (Figure 4a). Subsequently, we followed standard library construction steps, attaching sequencing adapters to both ends of the sequences before sequencing. The sequencing results were then decoded and analyzed to obtain the retained file information, with the deleted information being unrecoverable.

**Fig. 4.**
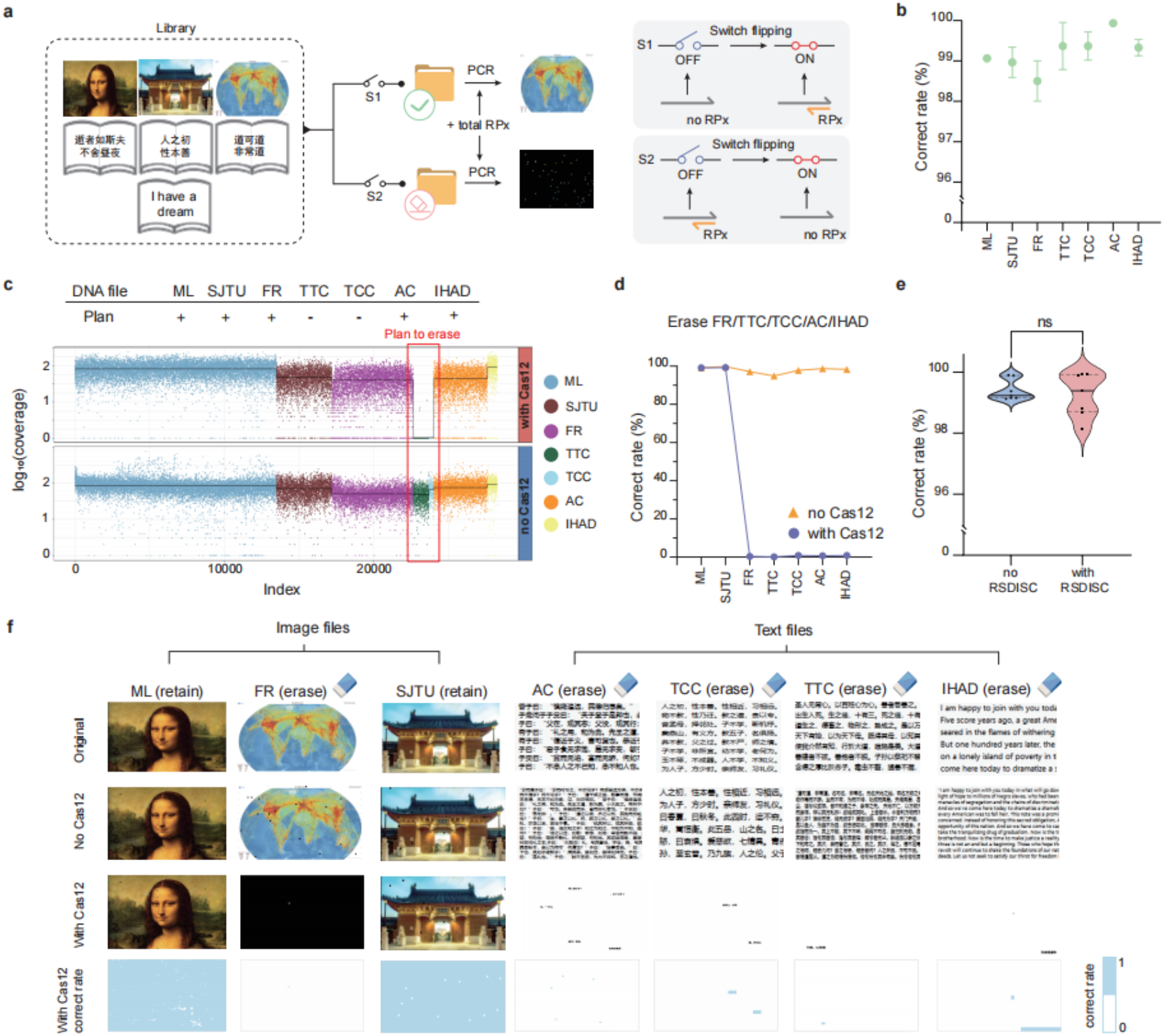
The application of the RSDISC system in files random sanitization. We encoded files of multimodal data into DNA templates based on the encoding principle (Supplementary Section). **a**, Schematic illustration of the selective erasure procedure. Typically, the RSDISC contains two types of starting switch: When both the primer and the template are present, switch S1 is closed, and data is retained; otherwise, switch S2 is closed and data is erased. **b**, Decoding accuracy when using the RSDISC assay to retain all 7 files. Error bars represent mean ± SD, where n = 3 replicates. **c**, Coverage comparison between with Cas12a-treated and no Cas12a-treated to erase TTC and TCC. The erasure efficiency for these two files exceeds 99%, underscoring the exceptional performance of RSDISC. **d**, Decoding accuracy comparison between with Cas12a-treated and no Cas12a-treated of erasing FR, TTC, TCC, AC and IHAD. **e**, Comparison of decoding accuracy of retaining all 7 files with RSDISC and without RSDISC to verify whether this system causes any loss of file data. With RSDISC means performing file erasure operations before library construction; no RSDISC means performing library construction directly. Data were analyzed by t-test. **f**, Reconstructed images and texts from NGS reads for the solutions after different treatments. The top-to-bottom presentation includes original files, results of without or with Cas12a treatment, and the distribution of correct rate.

To investigate whether this method would result in any loss of data from the targeted retained files, we added all the corresponding primers for the seven files before the erasure step to retain all of them. The results showed that the decoding accuracy for all seven files was above 98.5%, indicating that all the file information was essentially completely preserved (Figure 4b). Next, we chose to selectively erase the TTC and TCC files. The next-generation sequencing (NGS) results showed that the oligonucleotide sanitization efficiency for the TTC and TCC files reached 99.9%, indicating that the file sanitization efficiency of this method was 100% (SSIM < 0.1 is considered unrecoverable) (Figure 4c, Supplementary Figure 16). Then, we chose to retain the ML and SJTU files while erasing the other five files. The results showed that while achieving 100% file sanitization efficiency, the oligonucleotide sanitization specificity was as high as 99.5%, indicating that the file sanitization specificity as also 100% (Figure 4d). These results demonstrated that our method can efficiently erase non-target information while protecting the target information, and the erased information is unrecoverable.

To further verify whether the workflow of our method would affect the accuracy of the file data itself, we compared the decoding accuracy of the seven files prepared by Cas12a-treated library construction with that of conventional direct library construction. We found that the accuracy of both methods was essentially the same, indicating that our method does not cause additional data loss (Figure 4e). To further demonstrate the erasure effect of our method on files, we decoded the 7 files after different treatments and we found that our method has excellent erasure effects on file information (Figure 4f).

## Discussion

Convenient and reliable random permanent deletion of data stored in DNA is one of the fundamental challenges that must be addressed in DNA data storage, particularly for the secure preservation of big data^32-34^. In this work, we developed a method called RSDISC, which achieved selective erasure of files in a single step during storage, ensuring that the erased information cannot be recovered. Our method utilized the trans-cleavage activity of CRISPR-Cas12a, retaining the targeted oligonucleotides in the form of double-stranded structure and cleaving the residual ssDNA^35^. By applying our method to DNA storage pool containing 28,258 oligonucleotides, we demonstrated that the oligonucleotide sanitization efficiency up to 99.9% with oligonucleotide sanitization specificity up to 99.5%.

Sanitization with the RSDISC system relies on the characteristic of Cas12a, which can only non-specifically cleave ssDNA but not dsDNA^36^. In DNA storage, the templates encoding information are mainly generated through artificial synthesis technology and exist in the form of ssDNA oligonucleotides^37^. Therefore, we can use the trans-cleavage activity of Cas12a to selectively cleave single-stranded templates before library preparation to achieve the complete erasure of information-carrying DNA. Existing work mostly utilizes Cas12a to cleave ssDNA reporters less than 20 nucleotides long^38-40^. In contrast, we quantitatively explored the ability of Cas12a to trans-cleave long single strands (over 100 nucleotides) and verified that our system can efficiently cleave single strands with varying GC content (30% to 80%), different lengths (less than 150 nucleotides), and even those with severe secondary structure modifications. And the lengths of single strands investigated in this experiment match the length requirements for NGS^41^. Since Cas12a can trans-cleave single strands up to several thousand nucleotides long^20^, our technology has the potential for application on third-generation nanopore sequencing platforms^42,43^.

Compared to other random readout methods, the random erasure system we developed based on the trans-cleavage activity of CRISPR-Cas12a truly achieves complete information deletion, where the deleted sequences no longer exist in the storage system. This differs from existing random access deletion methods, such as selective deletion of data using strand displacement reactions or metastable hybridization-based random deletion systems^17-19,32^. oligonucleotides deleted using these methods are merely temporarily hidden within the system and can still be read later through direct sequencing or other means, making them unsuitable for information encryption. Our developed erasure method not only permanently and completely wipes information, ensuring it cannot be recovered by any means, but also achieves 100% efficiency and specificity in file erasure within the storage system. In a DNA pool consisting of 28,258 oligonucleotides, we achieved an oligonucleotide erasure efficiency of 99.9% and specificity of 99.5%. We also modeled the amplification efficiency and simulated that up to 10^11.7^ files can be erased, with a data capacity of up to 10^7^ GB. In the face of rapidly increasing data volumes in storage systems, selectively cleaving part of the templates before library preparation can effectively reduce non-specific hybridization between primers and templates in subsequent multi-step amplifications, thereby improving the accuracy of sequencing results. Additionally, our technology can also achieve selective data reading, significantly reducing the cost and time of sequencing by avoiding the need to sequence the entire database each time.

More generally, we believe that our work presents an important advance to the field of DNA information storage, with potential applications in information encryption, memory deallocation, file classification and so on. When migrating data or replacing storage devices, this method can be used to erase sensitive information, ensuring information security. Additionally, when updating data or dealing with insufficient storage capacity, our method allows for the thorough erasure of unnecessary information, freeing up memory for new data. Because our method possesses autonomous selectivity, we could construct different folders to selectively delete mixed information, thereby achieving separate storage of different types of data. With the rapid increase in storage data capacity, non-specific hybridization between templates and primers in complex systems becomes more severe. By cleaving a portion of templates, we can reduce non-specific hybridization caused by high-throughput oligonucleotides, further minimizing the impact on the accuracy of sequencing. Additionally, our single strands separation method can be applied in the field of molecular diagnostics to help reveal gene functions and regulatory mechanisms.

## Methods Materials

LbCas12a (1 μM) were purchased from New England Biolabs. Phusion High-Fidelity DNA Polymerase (2 U/μL), 5× HF buffer and dNTP (10 mM) were purchased from Thermo Fisher Scientific. PEG-200, Tris-HCl (pH 8.5, 10 mM), MgCl_2_ (1 μM), DTT (1 μM) were purchased from Sangon Biotech (Shanghai, China). The index sequences of the amplicons using an indexing kit (YEASEN). The VAHTS DNA Clean Beads was purchased from Vazyme Biotech (Nanjing, China). All RNA strands and DNA strands used in this study, except for oligonucleotide pools, were synthesized from Sangon Biotech (Shanghai, China). DNA primers were purified via high affinity purification (HAP), and other nucleic acid strands were purified via high-performance liquid chromatography (HPLC). The sequences of all the DNA strands and RNA strands that have been studied in this work, except for oligonucleotide pools, are summarized in Supplementary Tables 1. oligonucleotide pools were synthesized by Twist Biosciences (San Francisco, USA) and delivered in the form of DNA powder.

### Helper DNA preparation

The Helper DNA was produced by amplifying the ssDNA using a polymerase chain reaction in the CFX96 Touch Real-Time PCR Detection System (Bio-Rad Laboratories). In this procedure, 50 μL of the PCR contains 10 μL of 5× HF reaction buffer, 0.5 μL of DNA Polymerase (2 U/μL), 1 μL of dNTP (10 mM), 2 μL of MP6 ssDNA (10^5^ copies/μL), 2 μL of RP-Adapter (10 μM),2 μL of FP-Adapter/FP (10 μM) and 32.5 μL of nuclease-free water. The PCR was performed with the following protocol: (1) initial denaturation at 98 °C for 30 s, (2) denaturation at 98 °C for 10 s, (3) annealing at 63 °C for 30 s, (4) extension at 72°C for 30 s, with steps 2-4 repeated for 10 cycles, and then (5) final extension at 72 °C for 5 min. The PCR product was then purified from the reaction mixture using the magnetic beads, and the final elution volume is 20 μL.

### LbCas12a-crRNA complexes preparation

For the preparation of RNP complexes, 0.6 μL of LbCas12a (1 μM), 0.6 μL of crRNA ① (2 μM), 0.6 μL of crRNA② (2 μM), 1.0 μL of 10× Cas12 reaction buffer (10 mM Tris-HCl, pH 8.5, 10 mM NaCl, 15 mM MgCl_2_, 1 mM DTT), 0.5 μL of PEG-200 and 6.7 μL of nuclease-free water was incubated together at 37 °C for 30 min. The usage of each component can be adjusted according to the experimental consumption.

### Procedure of ssDNA selective protection assays

In a heterogeneous mixture of various single-stranded DNAs, selectively amplify the target-retained single strand using its corresponding reverse primer, and the ssDNA protection step can be combined with the Helper DNA preparation step. In a typical assay to protect x (x > 0) species of ssDNA, 10 μL of 5× HF reaction buffer, 0.5 μL of DNA Polymerase (2 U/μL), 1 μL of dNTP (10 mM), 2 μL of MP6 ssDNA (10^5^ copies/μL), 2 μL of RP-Adapter (10 μM),2 μL of FP-Adapter/FP (10 μM), 2 μL of ssDNA mixture (10^5^ copies/μL each), x types of RPs at a final reaction concentration of 400nM and nuclease-free water were mixed to total volume of 50 μL. The PCR was performed with the following protocol: (1) initial denaturation at 98 °C for 30 s, (2) denaturation at 98 °C for 10 s, (3) annealing at 63 °C for 30 s, (4) extension at 72°C for 30 s, with steps 2-4 repeated for 10 cycles, and then (5) final extension at 72 °C for 5 min. The PCR product was then purified from the reaction mixture using the magnetic beads, and the final elution volume is 20 μL.

### Procedure of ssDNA cleavage assay

In a simplex system, assays were performed containing 5 μL of 10× Cas12a reaction buffer, 19 μL of Helper DNA elution product, 2.5 μL of PEG-200, 2 μL of ssDNA (10^5^ copies/μL) and 21.5 μL of nuclease-free water. The reaction was initiated by adding 8 μL of LbCas12a-crRNA complexes with a final volume of 50 μL and incubated at 37 °C for 1 h. The product was then purified from the reaction mixture using the magnetic beads, and the final elution volume is 30 μL.

In a multiplex system, assays were performed containing 5 μL of 10× Cas12 reaction buffer, 19 μL of Helper DNA and ssDNA mixture elution product, 2.5 μL of PEG-200 and 23.5 μL of nuclease-free water. The reaction was initiated by adding 8 μL of LbCas12a-crRNA complexes with a final volume of 50 μL and incubated at 37 °C for 1 h. The product was then purified from the reaction mixture using the magnetic beads, and the final elution volume is 30 μL.

### DNA quantification via qPCR

In a typical qPCR protocol, we mixed 5 μL Blue Master Mix, 1 μL DNA product, the FPs and RPs were each at a final reaction concentration of 400nM and nuclease-free water in a reaction volume of 10μl. Blue SYBR Green Master Mix (YEASEN) was used for enzymatic amplification and fluorescence signal generation. The qPCR assays were performed on a CFX96 Touch Real-Time PCR Detection System using 96-well plates (Bio-Rad). Thermal cycling started with a 3-min incubation step at 95 °C, followed by 60 cycles of 10 s at 95 °C for DNA denaturing and 30 seconds at 60 °C for annealing and extension.

### NGS sequencing library preparation

First, depending on the combination of files to be accessed, the corresponding primers were pre-mixed to a final concentration of 4 μM. For each library, 19 μL of oligonucleotide pool after the selective cleavage and purification assay, 2.5 μL of FP-mix, 2.5 μL of RP-mix, 0.5 μL of polymerase, 1 μL of dNTP (10 mM), 10 μL of 5× HF buffer and 14.5 μL of nuclease-free water were mixed to total volume of 50 μL. The PCR was performed with the following protocol: (1) initial denaturation at 98 °C for 2 min, (2) denaturation at 98 °C for 20 s, (3) annealing at 63 °C for 30 s, (4) extension at 72°C for 30 s, with steps 2-4 repeated for 12 cycles, and then (5) final extension at 72 °C for 5 min. The product was then purified from the reaction mixture using the magnetic beads.

Next, 15 μL of purified amplicon, 2.5 μL of adapter FP-mix, 2.5 μL of adapter RP-mix, 0.5 μL of polymerase, 1 μL of dNTP (10 mM), 10 μL of 5× HF buffer and 18.5 μL of nuclease-free water were mixed to total volume of 50 μL. The PCR was performed with the following protocol: (1) initial denaturation at 98 °C for 2 min, (2) denaturation at 98 °C for 20 s, (3) annealing at 63 °C for 30 s, (4) extension at 72°C for 30 s, with steps 2–4 repeated for 3 cycles, and then (5) final extension at 72 °C for 5 min. The product was then purified from the reaction mixture using the magnetic beads. This step added the adaptor sequences to the ends of the amplicon.

The purified amplicon was then diluted 100-fold. Subsequently, the diluted amplicons were used as a template to perform a typical qPCR assay. The qPCR assays were performed containing 5 μL of Blue 2× Master Mix, 3 μL of diluted amplicon, 1 μL of N5 primer (diluted 5×) and 1 μL of N7 primer (diluted 5×). Thermal cycling started with a 3 min incubation step at 95 °C, followed by 30 cycles of 10 s at 95 °C for DNA denaturing and 30 s at 60 °C for annealing and extension. Ct values of the reactions were obtained by analyzing the qPCR results.

The index sequences were added to both ends of the amplicons using an indexing kit. Specifically, 15 μL of diluted amplicon, 1 μL of N5 primer, 1 μL of N7 primer, 0.5 of μL polymerase, 1 μL of dNTP (10 mM), 10 μL of 5× HF buffer and 21.5 μL nuclease-free water were mixed together. Thermal cycling started with a 3 min incubation step at 95 °C, followed by Ct + 4 cycles of 10 seconds at 95 °C for DNA denaturing and 30 s at 60 °C for annealing and extension. Finally, based on the quantification results, all libraries were pooled together and sent for NGS sequencing.

### Fluorophore quencher (FQ)-labeled reporter assays

5′-Cy5- and 5′-VIC-labeled ssDNA reporters were synthesized by Sangon Biotech. The assay was performed in a 0.1-mL PCR tube with a total reaction volume of 30 μL containing 19 μL of Helper DNA elution product, 0.288 μL of Cas12a (1 μM), 0.288 μL of crRNA① (2 μM), 0.288 μL of crRNA② (2 μM), 3.0 μL of 10× Cas12 reaction buffer, 1.5 μL of PEG-200, 1 μL of ssDNA reporter (60 μM, 30 μM, 15 μM, 7.5 μM, 3.0 μM, 1.8 μM, 0.6 μM) and 4.636 μL of nuclease-free water. Then the reactions were incubated in a fluorescence plate reader for up to 200 min at 37°C with fluorescence measurements taken every 15 s (Cy5: λ_ex_: 640 nm; λ_em_: 664 nm; VIC: λ_ex_: 526 nm; λ_em_: 543 nm). The following equation was used for each sample: net fluorescence intensity = fluorescence intensity (corresponding time points) – fluorescence intensity (background fluorescence without Cas12a). To determine the kinetic parameters of reporter, we established a standard curve to determine the relationship between fluorescence intensity and the concentration of cleaved reporter. The initial velocity (V_0_) calculated by using linear fit was plotted against reporter concentration to determine the Michaelis–Menten constants using GraphPad Prism software.

