## Supplemental information for "Random Sanitization in DNA information storage using CRISPR-Cas12a"

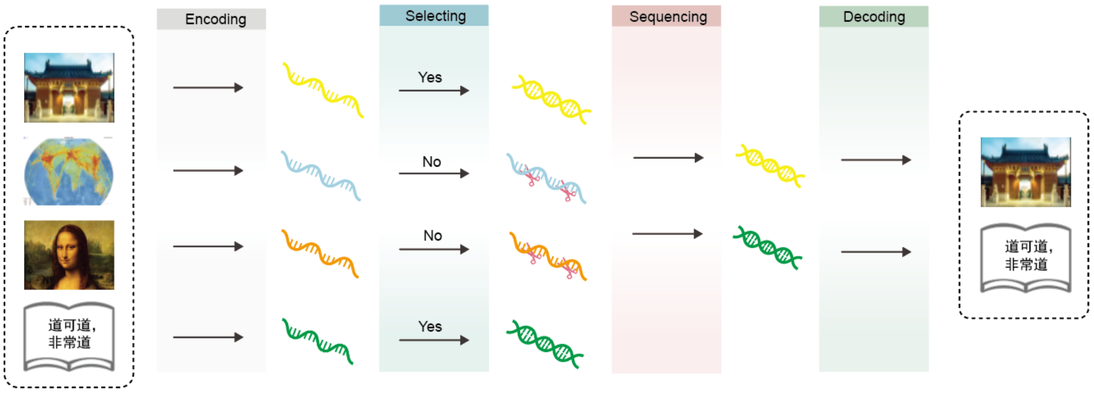


**Figure S1. Random sanitization system in DNA information storage overview.** The complete process of random sanitization method in DNA storage typically involves four segments: encoding, selecting, sequencing, and decoding. Initially, various types of materials, such as text and images are transformed into DNA sequences using specific coding scheme. Next, the primers encoding the target file DNA are used for amplification, and the trans-cleavage activity of Cas12a is employed for the selective deletion of file data. The retained DNA library is amplified and sequenced following the standard library preparation process. Detailed library preparation process refers to Methods. The utilization of fixed-energy primers ensures consistent amplification efficiency, leading to accurate decoding of individual files. Finally, decoding the NGS reads back into the bitmap image.

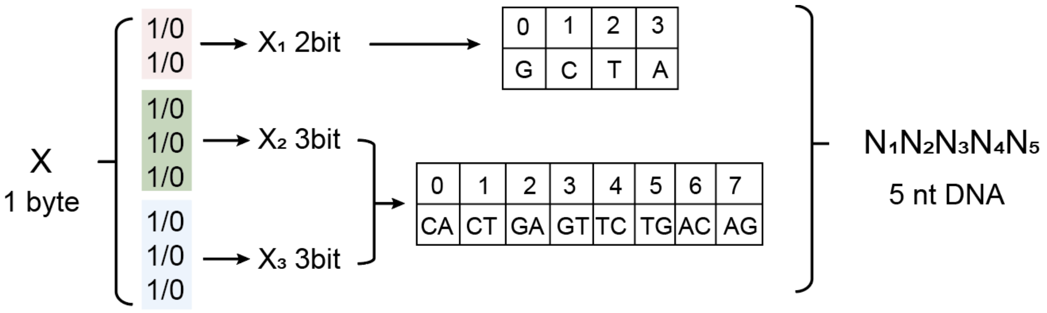


**Figure S2. Schematic illustrating the foundational logic of the encoding strategy.** Computer binary is converted into the quaternary representation of DNA bases using this encoding strategy.

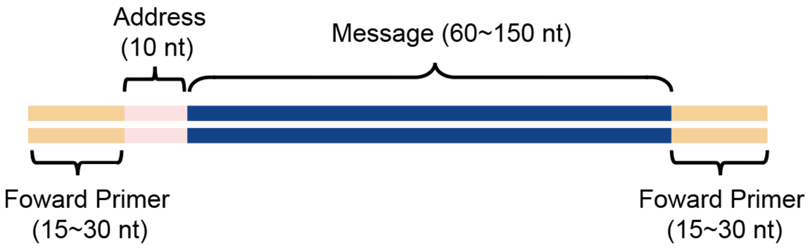


**Figure S3. Encoding a bitmap image or text in DNA sequence.** Given the synthesis limitations of Twist, the lengths of library primers, and the length of the truth marker binding site, we have roughly 70 nt per DNA oligonucleotide for the address and the message.

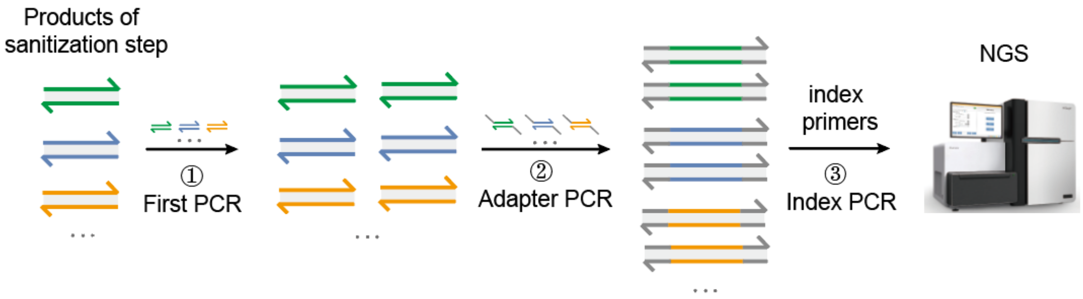


**Figure S4. Workflow of the library preparation before NGS.** This workflow involves of 3 steps of PCR using different types of primers. Firstly, we use normal primers to amplify the templates after step of ransom sanitization. Then, we add a common sequence (adapter) to both ends of the amplicons. After that, we use index primers to amplify the templates adding the index sequences to both ends of the amplicons. Finally, we send the prepared library for sequencing. Index primers were appended the index sequences of P5 and P7 to both ends of the normal primers.

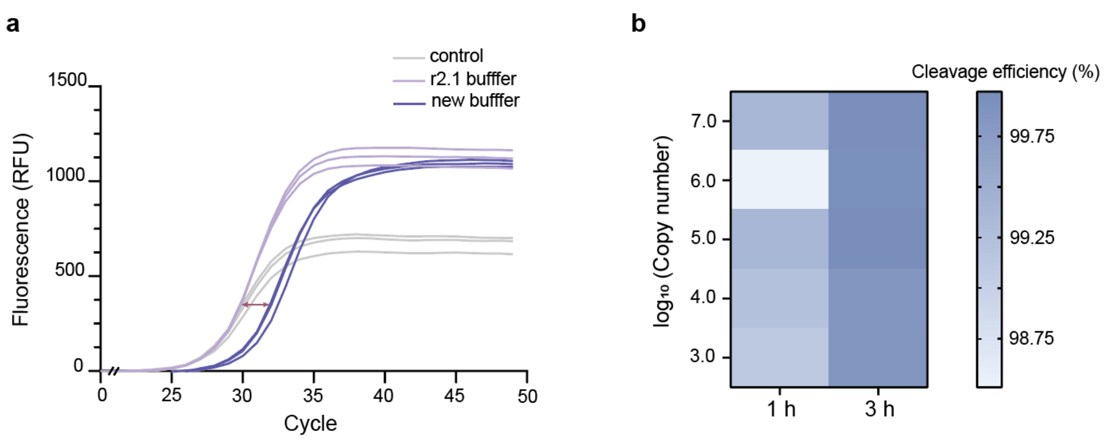


**Fig. S5 | Buffer and incubation time optimization for cleavage effect. a**, Time course assays of the ssDNA trans-cleavage efficiency of RSDISC assay with r2.1 buffer and new buffer. b, Showing the cleavage efficiency at different incubation times. The new buffer contains the following components: 10 mM Tris-HCl (pH 8.5), 10 mM NaCl, 15 mM $MgCl_2$, 1 mM DTT, 5% PEG-200.

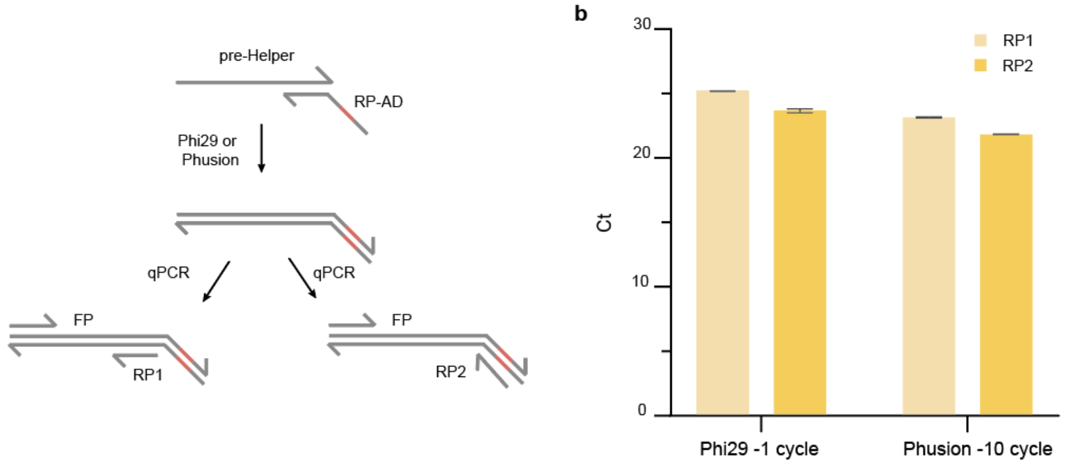


**Fig. S6 | The impact of different amplification methods on adapter ligation efficiency. a**, Schematic of comparing ligation efficiency using quantitative results by two primers. **b**, Quantitative results of pre-Helper substrate amplified using different polymerase enzymes.

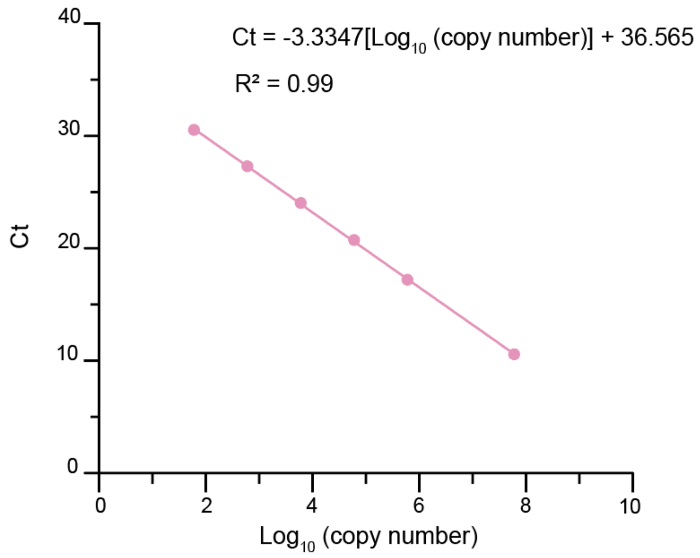


**Fig. S7 | Quantitative standard curve graph for 130 nt ssDNA (M7 template).** We will convert the Ct values obtained from qPCR measurements to copy numbers using the quantitative standard curve formula, and then calculate the cutting efficiency for each case. For cleavage efficiency mentioned in this work, the calculation formula is as follows: Cleavage efficiency (%) = 1- (copy number) $_{\text{with cas12a}}$ / (copy number) $_{\text{no cas12a}}$.

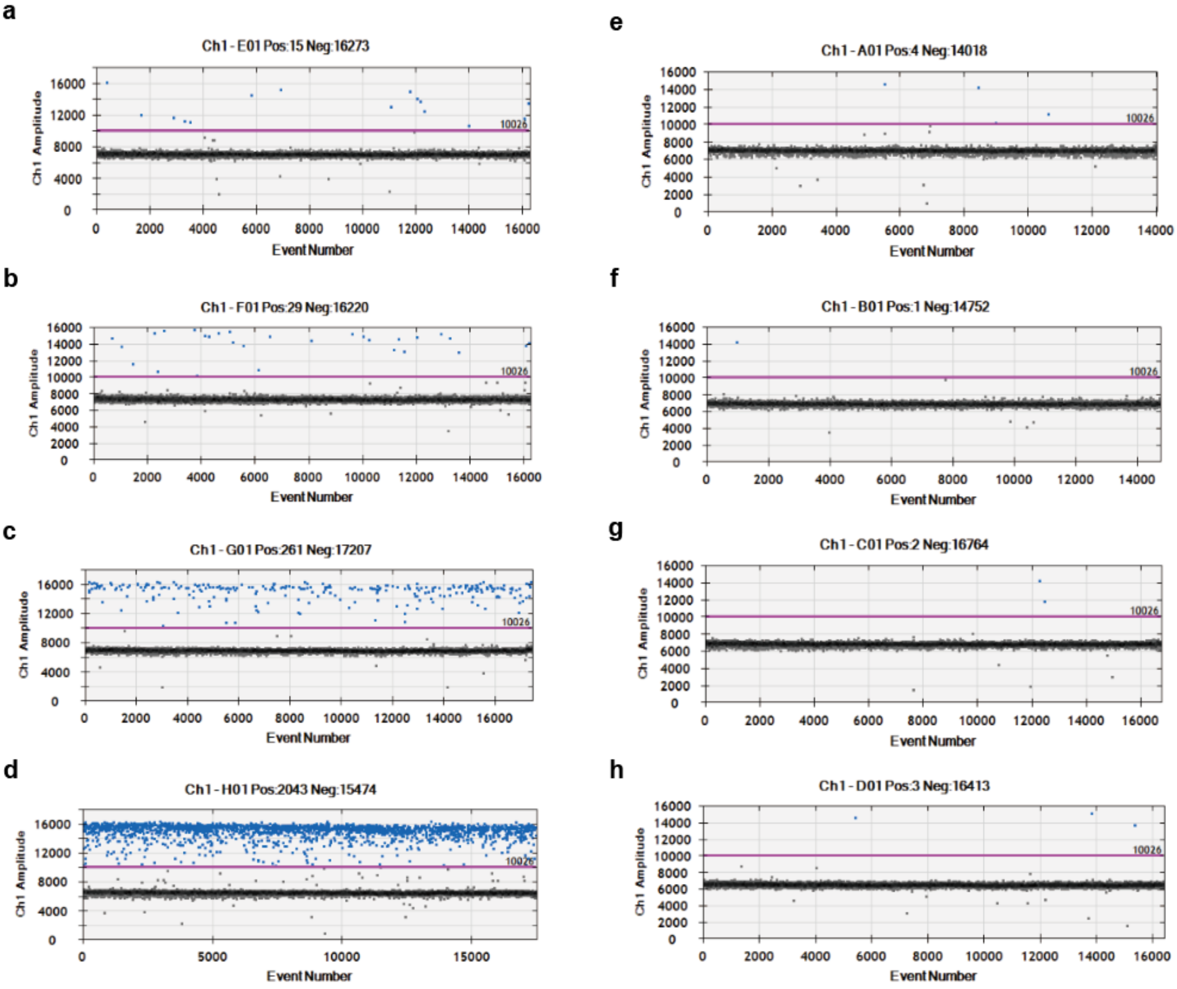


**Fig. S8 | Amplification efficiency diagram of ssDNA with/no Cas12a treatment by S-Helper system. a-d**, Amplification efficiency diagram with different initial copy numbers before Cas12a treatment by Droplet Digital $^{TM}$ PCR System. **e-h**, Amplification efficiency diagram with different initial copy numbers after Cas12a treatment by Droplet Digital $^{TM}$ PCR System.

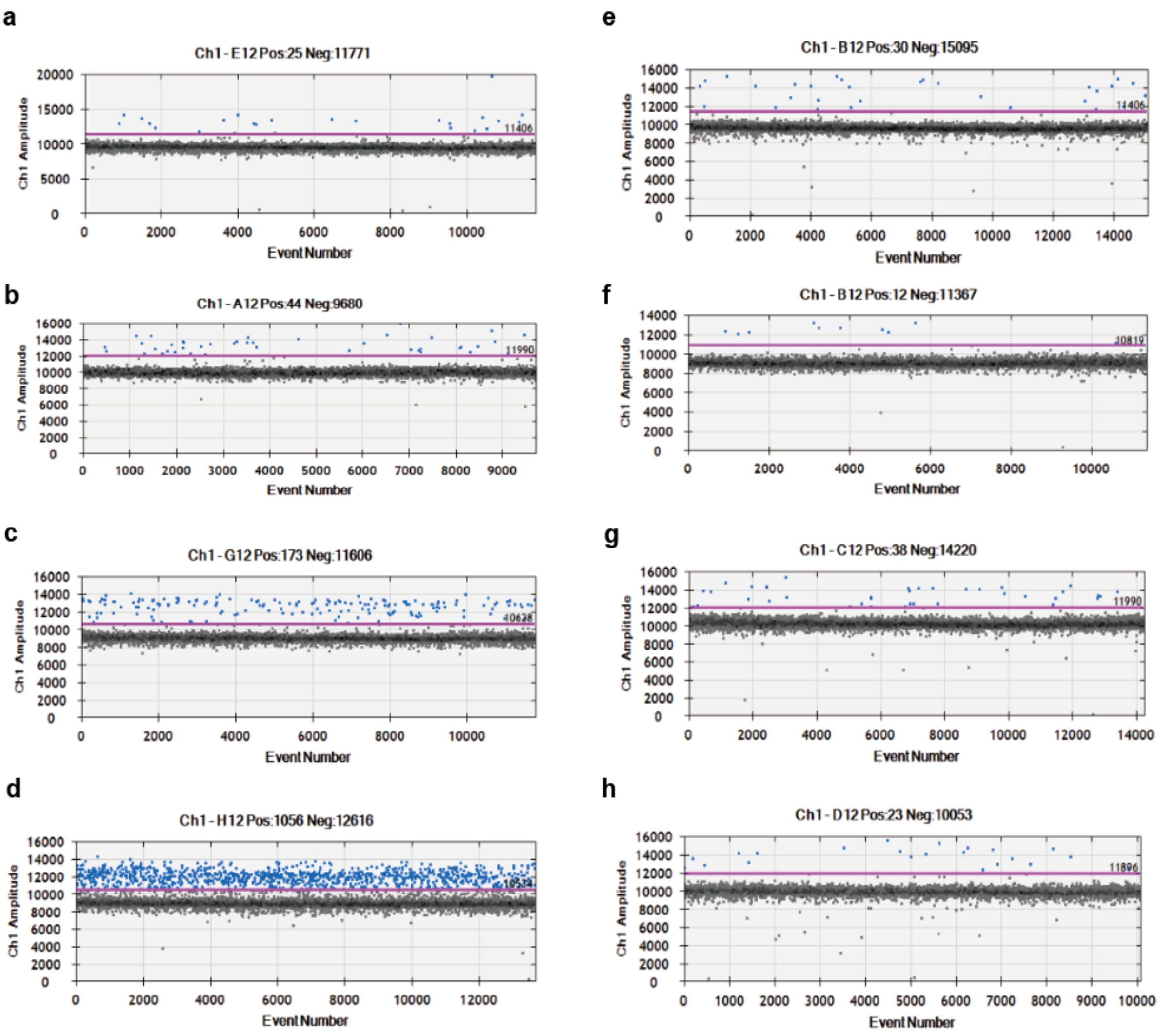


**Fig. S9 | Amplification efficiency diagram of ssDNA with/no Cas12a treatment by D-Helper system. a-d**, Amplification efficiency diagram with different initial copy numbers before Cas12a treatment by Droplet Digital ™ PCR System. **e-h**, Amplification efficiency diagram with different initial copy numbers after Cas12a treatment by Droplet Digital ™ PCR System.

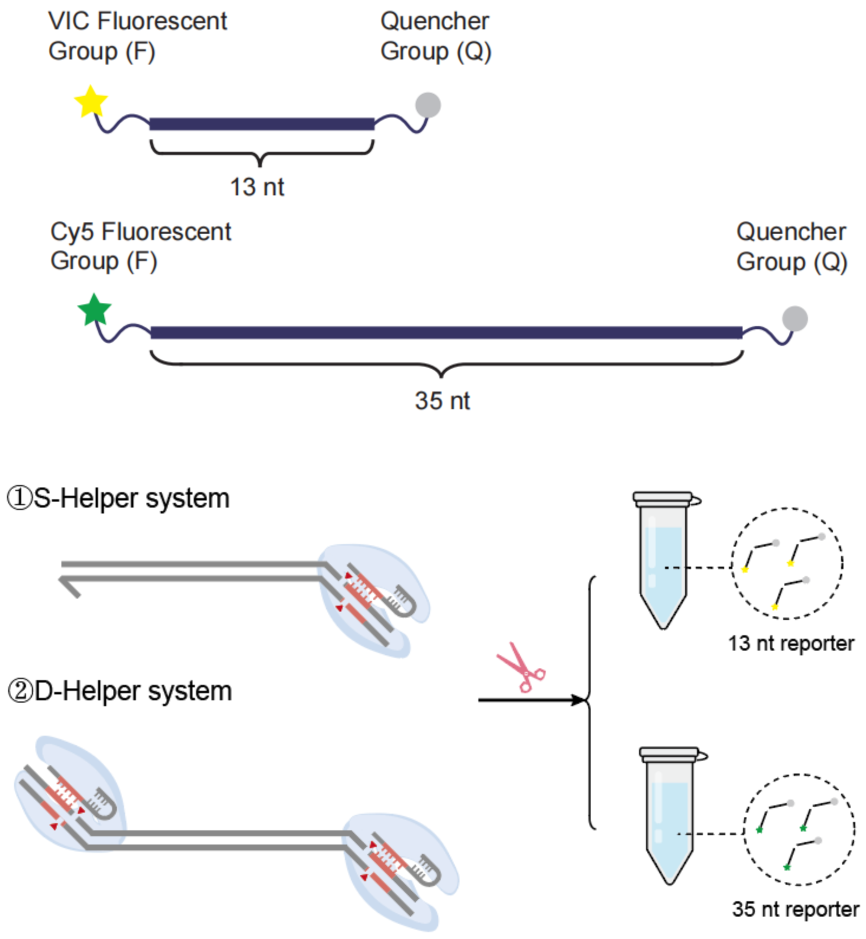


**Fig. S10 Kinetic detection method for the activation of Cas12 trans-cleavage activity in S-Helper and D-Helper systems.** To compare the efficiency of single-strand cleavage after activating Cas12a using S-Helper and D-Helper systems, we use the fluorometric method which is based on the generation of fluorescence signals by cleavage of FQ-labelled ssDNA reporters with the Cas12a trans-cleavage activities to define the cleavage efficiency.

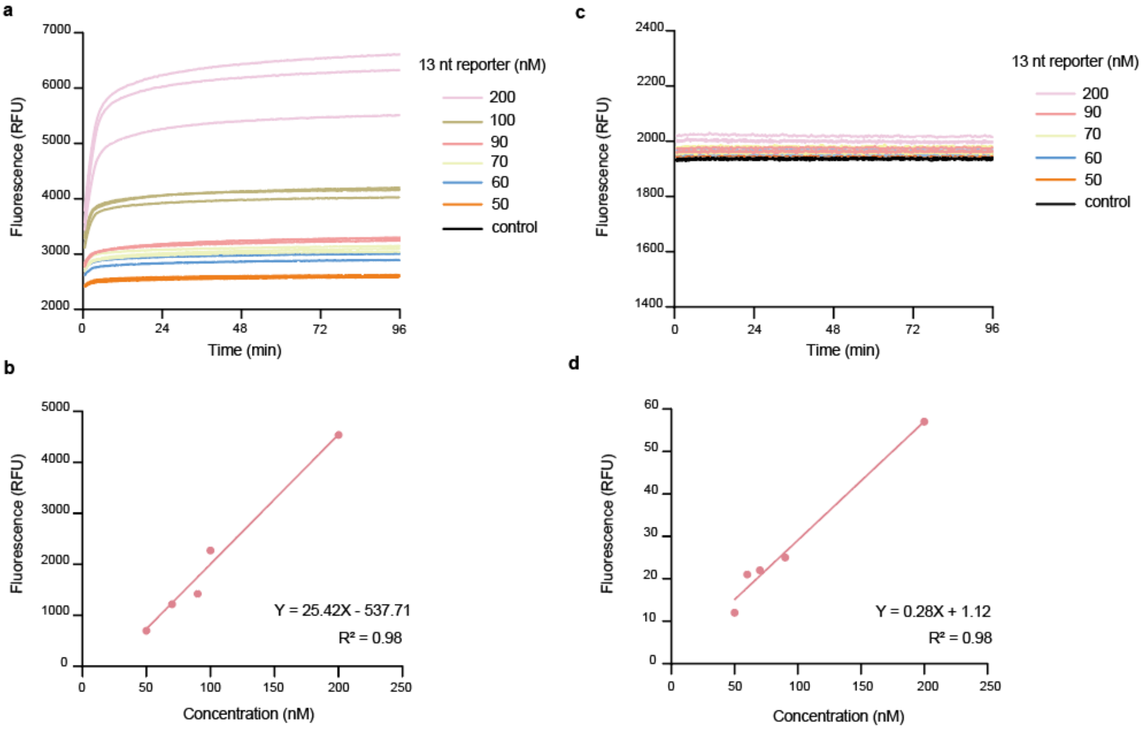


**Fig. S11 | Analysis of the Cas12a trans-cleavage kinetics for 13 nt reporter using a qPCR machine.** The trans-cleavage reaction systems were the same as shown in the Methods and all reactions were performed in a total volume of 20 μL with the fluorescence signals recorded by the a CFX96 Touch Real-Time PCR Detection System using 96-well plates (Bio-Rad). **a**, Fluorescence curves of Cas12a trans-reactions using different concentrations of the 13 nt ssDNA FQ-reporter. **b**, Fluorescence curves of the negative control experiments (NTC) corresponding to reaction systems shown in panel a. In NTC, the Cas12a enzyme was substituted with nuclease-free water, representing the un-cleaved signals. To establish a standard curve between the amounts of cleaved FQ-reporters and the fluorescence signals, all components but the FQ-reporter are premixed in the new buffer (specific components are detailed in Fig. S10). Reporter should be serially diluted, ranging from (0 ~ 200 nM), and then added to the reaction system to initiate the trans-cleavage reaction. **c**, Fluorescence curves of without Cas12a trans-reactions using different concentrations of the 13 nt ssDNA FQ-reporter. **d**, Calibrated curve with the background-subtracted fluorescence signals versus the concentrations of un-cleaved 13 nt reporters.

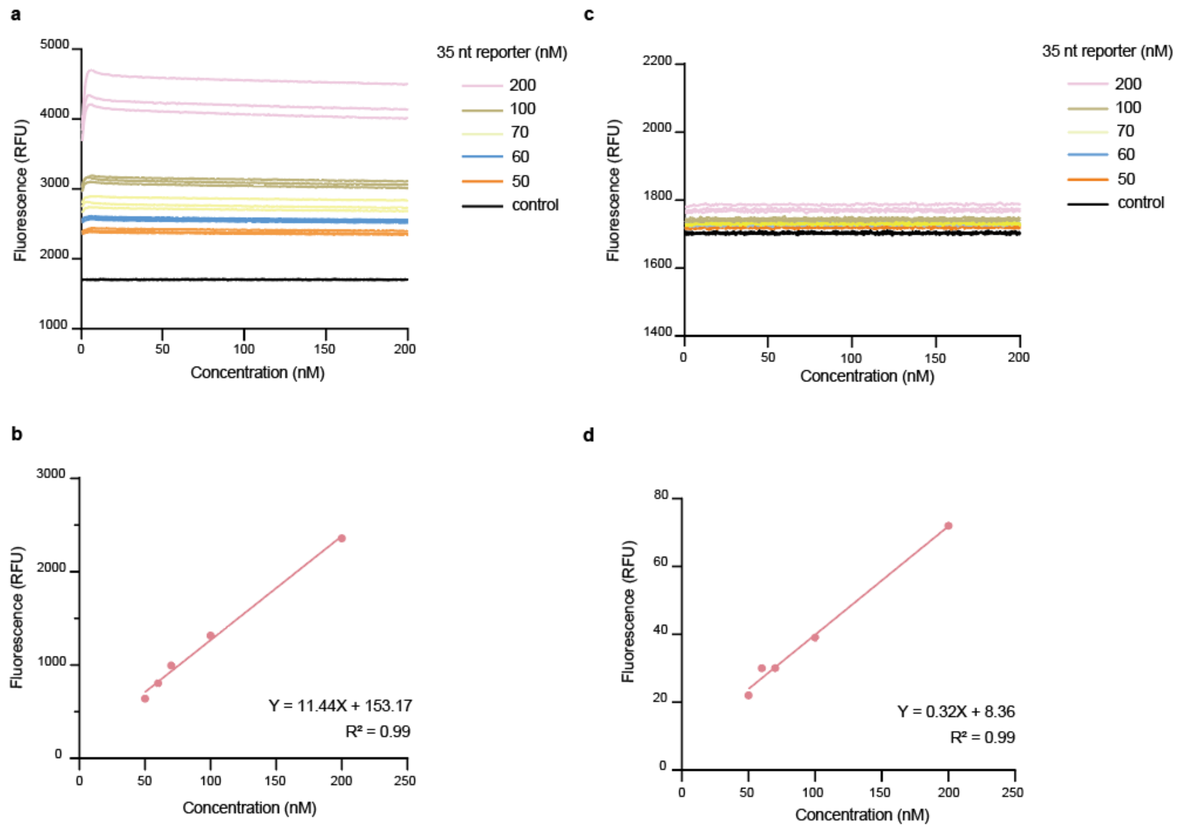


**Fig. S12 | Analysis of the Cas12a trans-cleavage kinetics for 35 nt reporter using a qPCR machine.** The trans-cleavage reaction systems were the same as shown in the Methods and all reactions were performed in a total volume of 20 μL with the fluorescence signals recorded by the a CFX96 Touch Real-Time PCR Detection System using 96-well plates (Bio-Rad). **a**, Fluorescence curves of Cas12a trans-reactions using different concentrations of the 35 nt ssDNA FQ-reporter. **b**, Fluorescence curves of the negative control experiments (NTC) corresponding to reaction systems shown in panel a. In NTC, the Cas12a enzyme was substituted with nuclease-free water, representing the un-cleaved signals. **c**, Fluorescence curves of without Cas12a trans-reactions using different concentrations of the 35 nt ssDNA FQ-reporter. **d**, Calibrated curve with the background-subtracted fluorescence signals versus the concentrations of un-cleaved 35 nt reporters.

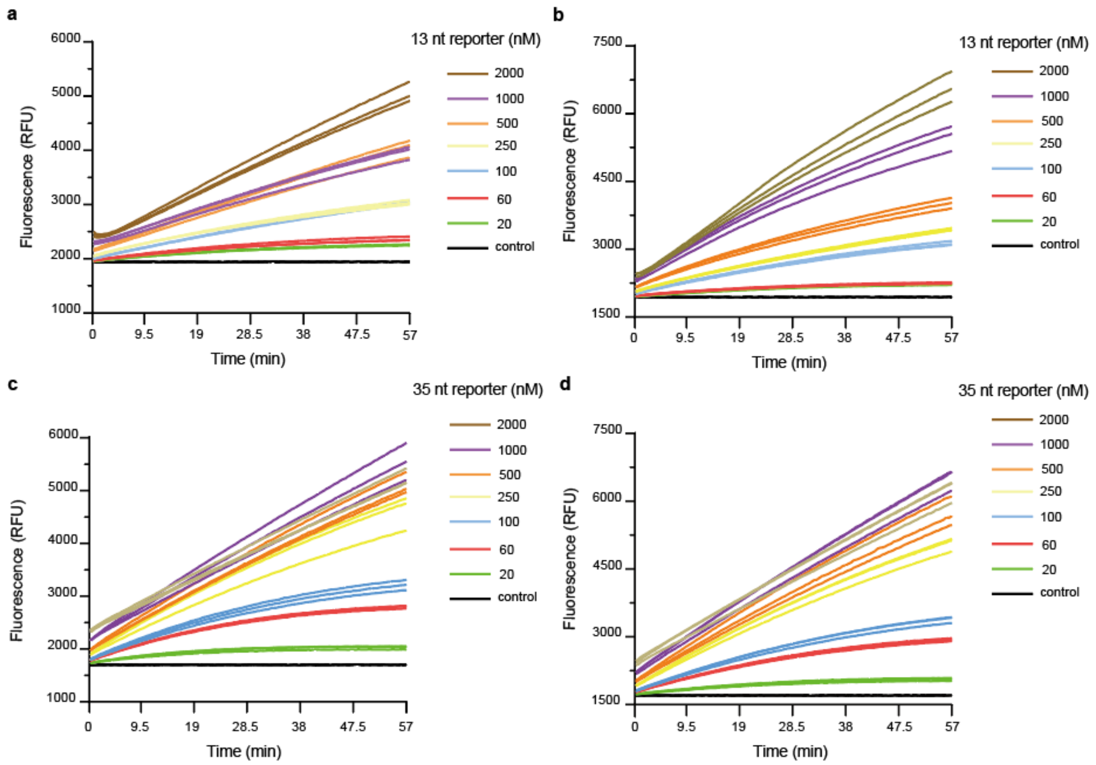


**Fig. S13 | Michaelis-Menten analysis of the Cas12a trans-cleavage kinetics in S-Helper and D-Helper systems for 13 nt or 35 nt reporters.** Representative plots of initial velocity versus time for a (**a**) 13 nt reporter with S-Helper-activated system, (**b**) 13 nt reporter with D-Helper-activated system, (**c**) 35 nt reporter with S-Helper-activated system, (**d**) 35 nt reporter with D-Helper-activated system.

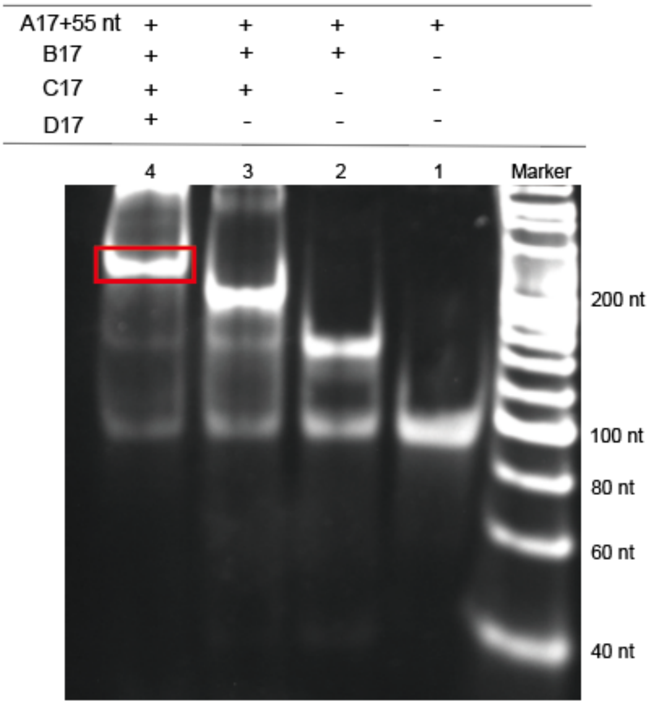


**Fig. S14 | The PAGE image of DTF.** Marker: DNA 20 bp marker; The DTF was assembled through mixing four DNA fragments of 17 nucleotides (A17+55 nt ssDNA, B17, C17, D17).

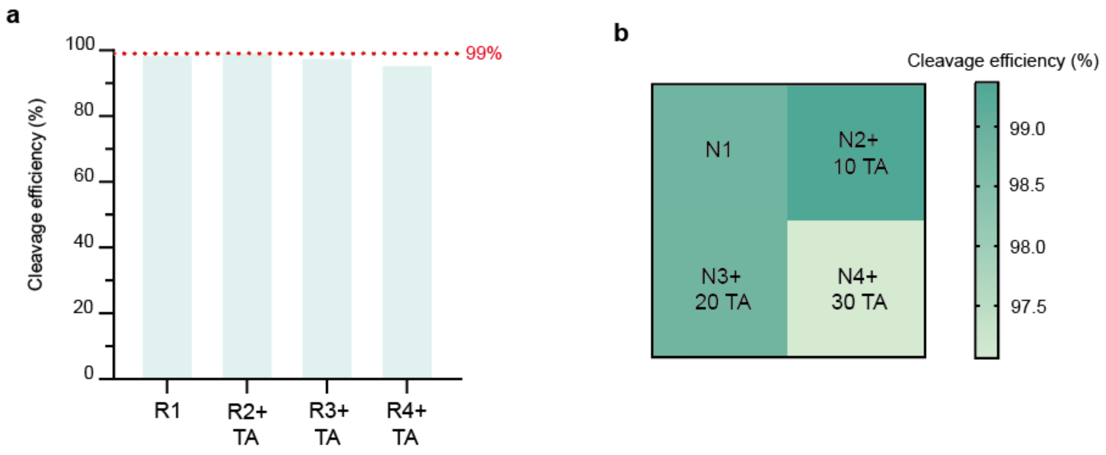


**Fig. S15 | The effect of T/A bases modification on single-strand cleavage efficiency. a,** The cleavage efficiency of single-strands with T/A bases modifications in a multiplex system. **b,** The effect of T/A bases modifications of different lengths on the same single-strand cleavage efficiency.

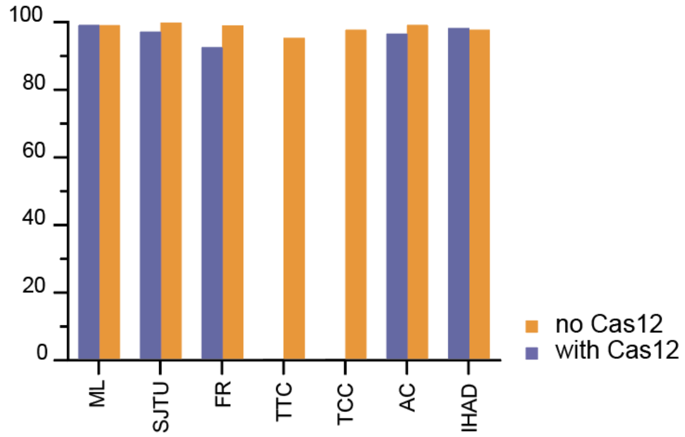


**Fig. S16 | The impact of selectively erasing TTC and TCC files on decoding accuracy.**

### Supplementary Table S1

| name | GC% | sequence |
|---|---|---|
| crRNA① | \ | UAAUUUCUACUAAGUGUAGAUUACCGAACGCUAAUAUCUGA |
| crRNA② | \ | UAAUUUCUACUAAGUGUAGAUAUCAACGCCCUAUAAGUCAA |
| M1 | 38% | CACTTTATCAGACACAGTTATGTGCTTGGGAATGAATGAGATTTCCCATCAACGGGAACAGGAGTTGGATGAGTACAGGTTTAGATGTAAATCTAGGCATCGTTATTAAAGTCGGTATAGTAATTTTTGC |
| M2 | 44% | GGATTCCCTAAGCTCTTCAATATTGCGACTTCACCTCAGAAAATAGGGATGTTAAGGGGTACCATCTAGCCGACCATGACAGTTCCGGGATTAATAGCTCCTGTGTGCTGACCATAATAGAACTACATTT |
| M3 | 58% | CCTAGCACCAGTTCTGCCTCCACCAGCAAGAACGACCCTGATTCTTGGAGACTAAGGTGTCTGTGGTCCAGTAATCCCAGCACTCTGGTCTGGGACGGAGAACGACCACCTTAGACGTCCCGTTCCCGGC |
| M4 | 66% | GATTCCCCCTCAGTCTCTCTGCCTCTCGGAAGGTCCTCACGACTTCACCTCGGATCAGCGGCTTGCCTCTCGGCCCGCACCACCTCGAGACCGTCGGCCTTGCAGGCGCACGCTCTGCCGCCTTCGTTCG |
| M4 + 3’ TA | 61% | GATTCCCCCTCAGTCTCTCTGCCTCTCGGAAGGTCCTCACGACTTCACCTCGGATCAGCGGCTTGCCTCTCGGCCCGCACCACCTCGAGACCGTCGGCCTTGCAGGCGCACGCTCTGCCGCCTTCGTTCGTTAATAATTT |
| M4 + 5’ TA | 61% | TTAATAATTTGATTCCCCCTCAGTCTCTCTGCCTCTCGGAAGGTCCTCACGACTTCACCTCGGATCAGCGGCTTGCCTCTCGGCCCGCACCACCTCGAGACCGTCGGCCTTGCAGGCGCACGCTCTGCCGCCTTCGTTCG |
| M5 | 53% | GGTGATGTGTGGATCTGCATGGTAAGCTGTCTGTCAGTACGCAGGGAATCAGAGC |
| M6 | 52% | TCTTCCCTATGGATGTGTTACAATGGATCAGGTACTGGTGGTCCTTCAACTCAGTTCCCATCGACCCGGAGGCTTACTTGGACTCTGCTTCGGCT |
| M7 | 48% | CCTAACACCAGTTCTTCCTCCACCAGCAAGAAAGACCAACCCAAGTACAATGACAATATAAGAAAGAGGTGGCTCATGCCTGTAATCCCAGCACTTTGGCCCATAGCTCTAGGTGGATCACTTGAGCCCA |
| S1 | 35% | ACCCAGGTGAGTTTTGTTTCACATAGCTTTAGTATATTCTTTATAGAACTGACAAAATTAGC |

| | | |
|---|---|---|
| | | CAGGCGGCATAATTGAATTCATTGGAACCCATTGACTAGATTAGTGAATTTGTGTGTATGT GGTTTCA |
| S2 | 36% | TCAGATGCTTTAGGCTCATGAGTTAACCTTAGTGGACTTCAAAACAGAGTTTATAACTACC AAGGAATACATTCCTTTAAGTCAAGAGAAATATCACCTTCCTAGAGTCATTAGTTTGTGAG TTGCTGTA |
| S3 | 39% | TCCGCAAAACCTACAATCTCTGAATCTTGGGAACTGAATACTGTGGTTACCTCAATGACAG TGGCTATACTGTAGTATTGTAGGAGGGAAATAACTCCCCAAGGGCATATAAACAAGAACA TTATCTTTT |
| S4 | 53% | CCTAGCACCAGTTCTGCCTCCACCAGCAAGAACGACCCTGATTCTTGGAGACTAAGGTGTC TGTGGTCCAGTAATCCCAGCACTCTGGTCTGGGACGGAGAACGACCACCTTAGACGTCCCG TTCCCGGCTTAATAATTT |
| S5 | 61% | GATTCCCCCTCAGTCTCTCTGCCTCTCGGAAGGTCCTCACGACTTCACCTCGGATCAGCGGC TTGCCTCTCGGCCCGCACCACCTCGAGACCGTCGGCCTTGCAGGCGCACGCTCTGCCGCCT TCGTTCGTTAATAATTT |
| seq1 | 58% | CCTAGCACCAGTTCTGCCTCCACCAGCAAGAACGACCCTGATTCTTGGAGACTAAGGTGTC TGTGGTCCAGTAATCCCAGCACTCTGGTCTGGGACGGAGAACGACCACCTTAGACGTCCCG TTCCCGGC |
| seq2 | 35% | ACCCAGGTGAGTTTTGTTTCACATAGCTTTAGTATATTCTTTATAGAACTGACAAAATTAGC CAGGCGGCATAATTGAATTCATTGGAACCCATTGACTAGATTAGTGAATTTGTGTGTATGT GGTTTCA |
| seq3 | 66% | GATTCCCCCTCAGTCTCTCTGCCTCTCGGAAGGTCCTCACGACTTCACCTCGGATCAGCGGC TTGCCTCTCGGCCCGCACCACCTCGAGACCGTCGGCCTTGCAGGCGCACGCTCTGCCGCCT TCGTTCG |
| seq4 | 36% | TCAGATGCTTTAGGCTCATGAGTTAACCTTAGTGGACTTCAAAACAGAGTTTATAACTACC AAGGAATACATTCCTTTAAGTCAAGAGAAATATCACCTTCCTAGAGTCATTAGTTTGTGAG TTGCTGTA |
| seq5 | 39% | TCCGCAAAACCTACAATCTCTGAATCTTGGGAACTGAATACTGTGGTTACCTCAATGACAG TGGCTATACTGTAGTATTGTAGGAGGGAAATAACTCCCCAAGGGCATATAAACAAGAACA TTATCTTTT |

| | | |
|---|---|---|
| R1 | 49% | ACTTCTGCCAACATTCAAATTCAGGTCTAGCCGTCCCAATAGAGGGTACCATCTTTGGAGC TAGACTGAGCCGTGACAAATGAACAAAAAAGCCATATCCAGGTTGCCCACCAGTGCCTCC AGCTTGCAC |
| R2 + TA | 50% | CTACTTGTACTCCAAAGTATCCTCGTCGTCCTACCTAATCTACTGGCCGCCTTAGCCAGGCG TGGACCTCGTGATCTGCCCGCCACCTAACATCATGGCAAGTCACTTGTCCTCCGAAGTTTA ATAATTT |
| R3 + TA | 60% | GATCAGCCCTCAGTCTCTCTGCCTCTCGGAAGGTCCTCACGACTTCACCTCGGATCAGCGG CTTGCCTCTCGGACCGACCTCACGAGCCGACTACAGGCGCGTAGCTGGGACTGCAGGCGCT TAATAATTT |
| R4 + TA | 70% | CGGCTCTGGGACGGCCTCCGTCTGGGCCGCACGCTCTGCCTCCCTCCCTCGTCTCGCCTGGT CCAGCCCCTCCCCCGCACCACCTCGAGACCGTCGGCCTGCTGCGCTCCGCTCTCGGTCTTA ATAATTT |
| N1 | 50% | ACTTCTGCCAACATTCAAATTCAGGTCTAGCCGTCCCAATAGAGGGTACCATCTTTGGAGC TAGACTGAGCCGTGACAAATGAACAAAAAAGCCATATCCAGGTTGCCCACCAGTGCCTCC AGCTTGCAC |
| N2 + 10TA | 50% | CTACTTGTACTCCAAAGTATCCTCGTCGTCCTACCTAATCTACTGGCCGCCTTAGCCAGGCG TGGACCTCGTGATCTGCCCGCCACCTAACATCATGGCAAGTCACTTGTCCTCCGAAGTTTA ATAATTT |
| N3 + 20TA | 50% | CTACTTGTACTCCAAAGTATCCTCGTCGTCCTACCCTACTGGCCGCCGCCAGGCGTGGACC TCGTGATCTGCCCGCCACCCATCATGGCAAGTCACTTGTCCTCCGAAGTTTAATAATTTTTA ATAATTT |
| N4 + 30TA | 50% | CTACTTGTACTCCCGTCGTCCTACCCTACTGGCCGCCGCCAGGCGTGGACCTCGTGATCTGC CCGCCACCCGGCATGGCAAGTCACTTGTCCTCCGAAGTTTAATAATTTTTAATAATTTTTAA TAATTT |
| 13 nt reporter | \ | /VIC/ AGGCAGCCGAAGG /MGB/ |
| 35 nt reporter | \ | /Cy5/ AATTCAGTTTCCTTCAAGATCCTCAAGAGAGCTTG /BHO3/ |
| RP-Adapter① | \ | CCATTTCTACCGAACGCTAATATCTGAGTCTCGTCTCGTGGGCTCGGAGATGTGTGGATTC CCTAAGCTCTTCAATATTGC |
| RP-Adapter② | \ | CCATTTCATCAACGCCCTATAAGTCAAGTCTCGTGGGCTCGGAGATGTGTGCAAAAATTAC TATACCGACTTTAATAACGA |

| | | |
|---|---|---|
| FP-Adapter① | \ | CCATTTCTACCGAACGCTAATATCTGAGTCTCGTCTCGTGGGCTCGGAGATGTGTCACTTTATCAGACACAGTTATGTGCT |
| FP-Adapter② | \ | CCATTTCATCAACGCCCTATAAGTCAAGTCTCGTGGGCTCGGAGATGTGTCACTTTATCAGACACAGTTATGTGCT |
| ML-FP | \ | ACCAATGGGAGTCACTGCTG |
| ML-RP | \ | CGTACACTGGATCAGCGTCG |
| ML-FP-AD | \ | TCGTCGGCAGCGTCAGATGTGTATAAGAGACAGACCAATGGGAGTCACTGCTG |
| ML-RP-AD | \ | GTCTCGTGGGCTCGGAGATGTGTATAAGAGACAGCGTACACTGGATCAGCGTCG |
| SJTU-FP | \ | GCTCTTCCTCTCACATCTTTATTTAACC |
| SJTU-RP | \ | CACTGCCAGCTTGTGCCT |
| SJTU-FP-AD | \ | TCGTCGGCAGCGTCAGATGTGTATAAGAGACAGGCTCTTCCTCTCACATCTTTATTTAACC |
| SJTU-RP-AD | \ | GTCTCGTGGGCTCGGAGATGTGTATAAGAGACAGCACTGCCAGCTTGTGCCT |
| FR-FP | \ | GGATGGGACTCCAATGCAAAACT |
| FR -RP | \ | GAAGCCAGATCTCAAAGTGTCCT |
| FR -FP-AD | \ | TCGTCGGCAGCGTCAGATGTGTATAAGAGACAGGGATGGGACTCCAATGCAAAACT |
| FR -RP-AD | \ | GTCTCGTGGGCTCGGAGATGTGTATAAGAGACAGGAAGCCAGATCTCAAAGTGTCCT |
| TTC-FP | \ | TGAAAGACGTCACAGCAAGGT |
| TTC -RP | \ | GCTGAGCTGTCACATCACTTCA |
| TTC -FP-AD | \ | TCGTCGGCAGCGTCAGATGTGTATAAGAGACAGTGAAAGACGTCACAGCAAGGT |
| TTC -RP-AD | \ | GTCTCGTGGGCTCGGAGATGTGTATAAGAGACAGGCTGAGCTGTCACATCACTTCA |
| TCC-FP | \ | TGTAGGAGAGATTGGGCTAGAGAG |
| TCC -RP | \ | CCACACTCTGCCTCTCATGGTAT |
| TCC -FP-AD | \ | TCGTCGGCAGCGTCAGATGTGTATAAGAGACAGTGTAGGAGAGATTGGGCTAGAGAG |
| TCC -RP-AD | \ | GTCTCGTGGGCTCGGAGATGTGTATAAGAGACAGCCACACTCTGCCTCTCATGGTAT |
| AC-FP | \ | GGATGGGACTCCAATGCAAAACT |
| AC -RP | \ | AAGTTACTGAAGGATATGCCACATCA |
| AC -FP-AD | \ | TCGTCGGCAGCGTCAGATGTGTATAAGAGACAGGGATGGGACTCCAATGCAAAACT |
| AC -RP-AD | \ | GTCTCGTGGGCTCGGAGATGTGTATAAGAGACAGAAGTTACTGAAGGATATGCCACATCA |
| IHAD-FP | \ | CCTCATCTGTAAAGCAGGGAGAGA |

| | | |
|---|---|---|
| IHAD -RP | \ | ACAGCCATCAGATATCCAGCAG |
| IHAD -FP-AD | \ | TCGTCGGCAGCGTCAGATGTGTATAAGAGACAGCCTCATCTGTAAAGCAGGGAGAGA |
| IHAD -RP-AD | \ | GTCTCGTGGGCTCGGAGATGTGTATAAGAGACAGACAGCCATCAGATATCCAGCAG |

**Supplementary Table S2**. Comparison of the Cas12a kinetic parameters from different research groups

| Research groups | $K_m$ (M) | $K_{cat}/K_m$ ($M^{-1}$ $s^{-1}$) | Ref. |
|---|---|---|---|
| **Chen *et al.*** | $7.25 \times 10^7$ | $1.7 \times 10^7$ | (Chen et al., 2018) |
| **Ramachandran *et al.*** | $2.13 \times 10^{-7}$ | $4.22 \times 10^5$ | (Ramachandran and Santiago, 2021) |
| **Shen *et al.*** | $1.43 \times 10^{-7}$ (S-Helper system)<br>$1.85 \times 10^{-7}$ (D-Helper system) | $8.44 \times 10^4$ (S-Helper system)<br>$8.49 \times 10^4$ (D-Helper system) | This work |
